# Single nucleus RNASeq profiling of mouse lung: reduced dissociation bias and improved detection of rare cell types compared with single cell RNASeq

**DOI:** 10.1101/2020.03.06.981407

**Authors:** Jeffrey R Koenitzer, Haojia Wu, Jeffrey J Atkinson, Steven L Brody, Benjamin D Humphreys

## Abstract

**RATIONALE:** Single cell RNA-sequencing (scRNASeq) has led to multiple recent advances in our understanding of lung biology and pathophysiology, but utility is limited by the need for fresh samples, loss of cell types due to death or inadequate dissociation, and the induction of transcriptional stress responses during tissue digestion. Single nucleus RNASeq (snRNASeq) has addressed these deficiencies in some other tissues, but no protocol exists for lung. We sought to develop such a protocol and compare its results with scRNA-seq.

**METHODS:** Single nucleus suspensions were prepared rapidly (45 min) from two mouse lungs in lysis buffer on ice while a single cell suspension from an additional mouse lung was generated using a combination of enzymatic and mechanical dissociation (1.5 h). Cells and nuclei were processed using the 10x Genomics platform, and following sequencing of cDNA libraries single cell data was analyzed by Seurat.

**RESULTS:** 16,656 single nucleus and 11,934 single cell transcriptomes were generated. Despite reduced mRNA levels in nuclei vs. cells, gene detection rates were equivalent in snRNASeq and scRNASeq (∼1,750 genes and 3,000 UMI per cell) when mapping intronic and exonic reads. snRNASeq identified a much greater proportion of epithelial cells than scRNASeq (46% vs 2% of total), including basal and neuroendocrine cells, while reducing immune cells from 54% to 15%. snRNASeq transcripts are enriched for transcription factors and signaling proteins, with reduced detection of housekeeping genes, mitochondrial genes, and artifactual stress response genes. Both techniques improved mesenchymal cell detection over previous studies, and analysis of fibroblast diversity showed two transcriptionally distinct populations of *Col13a1+* cells, termed *Bmper+* and *Brinp1+* fibroblasts. To define homeostatic signaling relationships among cell types, receptor-ligand mapping of was performed for alveolar compartment cells using snRNASeq data, revealing complex interplay among epithelial, mesenchymal, and capillary endothelial cells.

**CONCLUSION:** Single nucleus RNASeq can be readily applied to snap frozen, archival murine lung samples, improves dissociation bias, eliminates artifactual gene expression and provides similar gene detection compared to scRNASeq.

## Introduction

The emergence of single cell RNA sequencing (scRNASeq) technologies in the last decade has led to a rapid phase of discovery in lung research, including the identification of ionocytes in airway epithelium and characterization of pro-fibrotic macrophages and aberrant basaloid cells in idiopathic pulmonary fibrosis^1,2,3^. An emerging alternative to scRNASeq is single nucleus RNASeq (snRNASeq), which generates transcriptomic information from isolated nuclei. This approach has previously been reported in brain^4,5,6^ and kidney^7^ and unlike scRNASeq can be readily applied to cryopreserved, archival samples.

Notably, there are additional potential advantages to snRNASeq. Depending on conditions and technique, dissociation protocols used to generate single cell suspensions in adult tissues can underrepresent fragile cell types or fail to liberate matrix-embedded mesenchymal cells. Indeed, single cell RNASeq data from prior studies in mouse and human lung have shown bias toward immune cell types with underrepresentation of airway/alveolar epithelial cells and fibroblasts^8,9^, populations which are key drivers of pathologies including fibrosis^10^. Additionally, snRNASeq reduces sequencing of housekeeping and mitochondrial genes in favor of cell identity relevant genes such as transcription factors and long noncoding RNA, which may improve cell type differentiation versus scRNASeq at a given read depth^6,7^. While mRNA in the nucleus are often incompletely processed and gene detection rates in snRNASeq are poor when mapped to exons alone, previous studies suggest performance is similar when intronic reads are included during alignment^6,7^.

Here, we modified an existing protocol for isolation of human lung nuclei developed for the Human Cell Atlas^11^ for cryopreserved mouse lung, such that a FACS purification step was unnecessary. In parallel, we generated single cell suspensions from healthy mouse lung. We compared results head to head in terms of sensitivity, cell representation, transcriptional stress responses and differential gene expression, and characterized fibroblast diversity and intercellular receptor-ligand signaling in our data.

## Methods

### Single cell preparation from fresh mouse lung

Single cell suspensions were obtained from the lungs of one wild type adult C57Bl/6J mouse by a combination of enzymatic and mechanical dissociation (GentleMacs, Miltenyibiotec), essentially as described^12^. To clear debris, RBCs, and dead cells, the resulting cell suspension was applied to an OptiPrep density gradient with 12%, 18%, and 30% layers and centrifuged for 15 min at 600x*g*. All cells above the RBC layer were removed, diluted in 50 ml PBS, centrifuged again at 1000x*g* for 20 min, washed in 50 ml PBS + 0.1% BSA, counted, and diluted to 10,000 cells/µl.

### Single nucleus preparation from frozen mouse lung

Lung tissue from two wild type adult C57Bl/6J mice was isolated at the time of sacrifice and snap frozen in liquid nitrogen. Nuclei were prepared from frozen tissue under RNAse-free conditions by a method adapted from an existing protocol^11^. Briefly, samples were cut to ∼7 mm pieces, injected (26G needle) with 1 mL ice-cold Nuclei EZ Lysis buffer (NUC-101, Sigma-Aldrich) supplemented with protease (Roche #589279100) and RNAse (Promega #N2615, Life Technologies #AM2696) inhibitors, and minced to 1-2 mm pieces with scissors in a weigh boat with 1 mL lysis buffer. The sample was then transferred to a GentleMacs C tube and an additional 1 mL lysis buffer was added. The GentleMacs™ *lung1* and *lung2* programs were run in sequence and the latter stopped after 20 s. Foam was spun down for 1 min at 750x*g*. The suspension was passed through a 40 µm cell strainer and washed with 4 mL cold PBS with 0.1% BSA and 0.1% RNase inhibitor, and then passed through a 5 µm filter (pluriSelect). Nuclei were pelleted at 600 x *g*, resuspended in 1xPBS with 0.1% BSA and 0.1% RNAse inhibitor, counted, and diluted to 10,000 nuclei/µl.

### Library preparation, sequencing, and bioinformatics

10x Chromium libraries were prepared according to manufacturer protocol (10x Genomics) and submitted for sequencing through the Washington University Genome Technology Access Center (GTAC) on a NovaSeq S4 flow cell. Raw sequencing data were processed using the zUMIs pipeline^13^. Briefly, low quality barcodes were removed using an internal zUMIs read filtering algorithm, followed by mapping of the remaining barcodes to the mouse genome (mm10) using STAR 2.5.3a, and expression matrices containing intronic, exonic, and intronic+exonic reads were generated for both scRNASeq and snRNASeq data.

Data were further processed using Seurat. For quality control, only genes expressed in >5 cells and cells expressing at least 200 genes were retained. No mitochondrial gene expression cutoff was used for single nucleus data, while cells expressing >10% mitochondrial genes were excluded from single cell analysis. Data were normalized and scaled, and the number of principal components estimated using *RunPCA* followed by *ElbowPlot*. Dimensionality reduction was performed using *RunUMAP*. Markers for cell clusters were identified using the *FindAllMarkers* function in *Seurat*, and cell types were annotated using canonical markers. To combine independent experiments, datasets were merged using the *Harmony* package to correct for batch effects. To further resolve immune cells, the *Subset* function in *Seurat* was used, with subsets of cells subjected to a second round of principal components identification and dimensionality reduction as above. Clusters of presumed doublets were identified and removed manually (relying on the presence of marker genes from multiple clusters and relatively high UMI counts).

### Merging of cell and nucleus datasets

All cells (11,473) and nuclei (16,656) were merged into a single object using the *merge* function followed by batch correction with *Harmony*. PCA identification, clustering, and cell type annotation were then performed by the same workflow as above. Mesenchymal cells were isolated using *subset* for fibroblast, pericyte, and smooth muscle cell clusters. For some analyses, *AverageExpression* was used to compare gene expression in all clusters rather than within clusters, otherwise, expression was profiled using the *DoHeatmap, Dotplot, Vlnplot*, and *FeaturePlot* functions within Seurat.

### Receptor-Ligand Analysis

To identify ligand-receptor interactions, we grouped cell types from the alveolar milieu and employed a curated ligand-receptor (LR) list with 2,557 LR pairs, as described^7^. Receptors and ligands were selected on the basis of their differential expression in the selected subgroups of alveolar cells. For plotting, ligands and receptors with unique expression (q-val > 0.75) were selected. The gplots package function *heatmap*.*2* was used for visualization.

### Data Availibility

The accession number for the RNA sequencing data reported in this paper is NCBI GEO: GSE145998

## Results

### snRNASeq and scRNASeq have similar gene detection rates and dissociation bias is reduced in snRNASeq

Based on previous experience in kidney nuclear isolation for snRNASeq in which prolonged isolation and washing steps resulted in RNA degradation and poor quality cDNA libraries, we avoided staining and flow sorting of nuclei which reduced total isolation time to 45 min. Given the incomplete processing of nuclear mRNA, we compared gene detection when mapping to exons, introns, or both in both snRNASeq and scRNASeq. Gene detection per nucleus in our data was similar to detection per cell as long as both intronic and exonic reads were included during genome mapping (Fig 1A). Surprisingly, and contrary to previous reports^6,7^, inclusion of intronic reads also improved gene detection in scRNASeq, though to a lesser extent (Fig 1A). Accordingly, we included intronic and exonic reads for all samples during sequence alignment.

**Figure 1.**
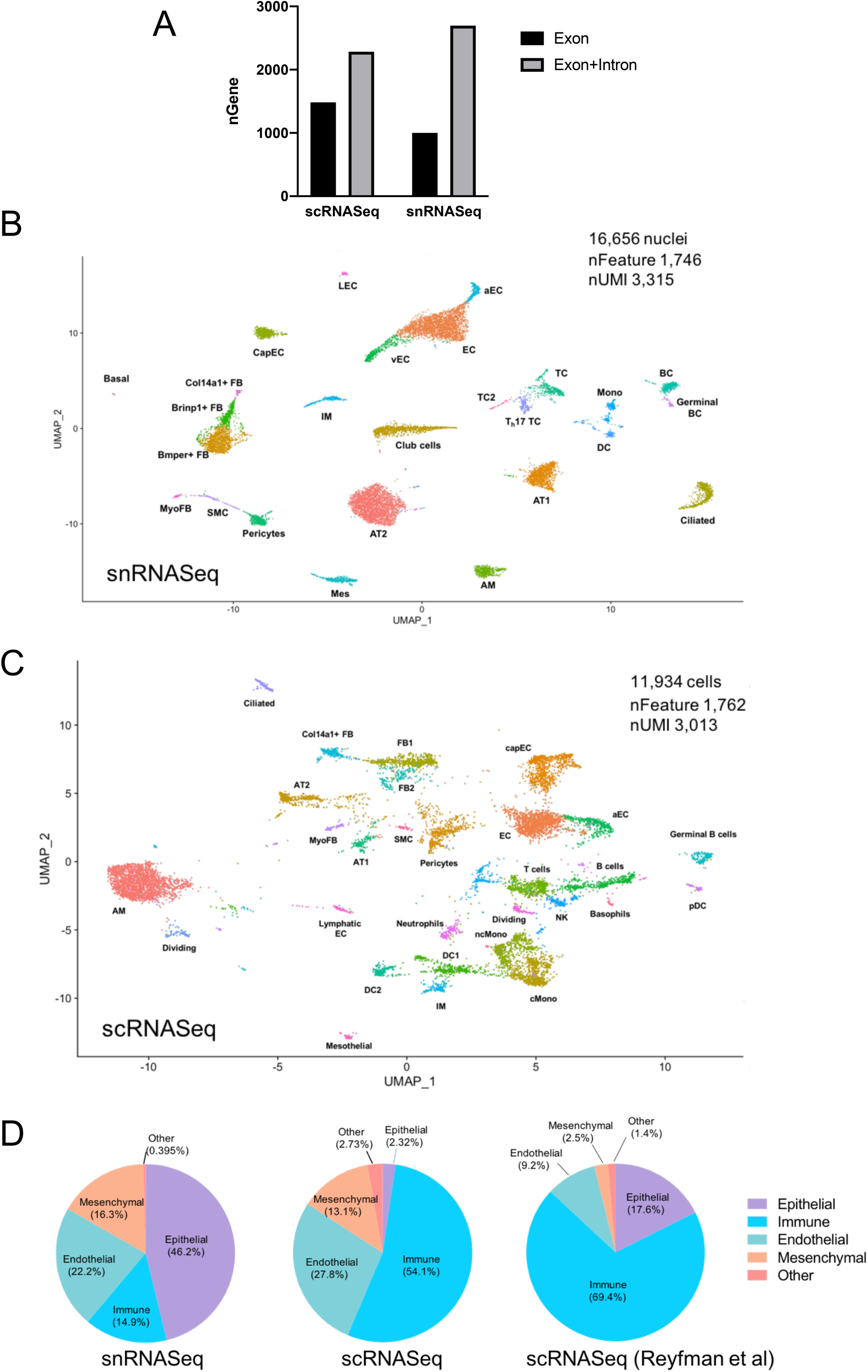
Single nucleus RNASeq offers similar gene detection and improves dissociation bias compared to scRNASeq. (A) Genes detected per cell when exonic reads alone or exonic and intronic reads are used in mapping. (B) Uniform manifold approximation and projection (uMAP) plot of 16,656 nuclei from snRNAseq data, combined from two replicates, with annotation of cell types. (C) Annotated uMAP plot of 11,473 cells from scRNASeq. (D) Percentages of total cells/nuclei grouped by category from snRNASeq, scRNASeq, and a previously published scRNASeq dataset. EC, endothelial cells; aEC, arterial endothelial cells; vEC, venous endothelial cells; CapEC, capillary endothelial cells; LEC, lymphatic endothelial cells; AT1, alveolar type 1 epithelial cells; AT2, alveolar type 2 epithelial cells; BC, B cells; TC, T cells; DC, dendritic cells; FB, fibroblasts; MyoFB, myofibroblasts; AM, alveolar macrophages; IM, interstitial macrophages; NK, natural killer cells; Mes, mesothelial cells; SMC, smooth muscle cells.

16,656 single nucleus libraries were generated from snap frozen lungs of two wild type mice, in parallel with 11,934 single cell libraries from one additional mouse. Unsupervised clustering of single nucleus data initially resulted in 25 clusters after dimensional reduction using Seurat v3 (Fig 1B). Alveolar and airway epithelial cell types were represented, including basal cells, neuroendocrine cells (in one dataset, Fig S1), and club cells, none of which were seen in our scRNASeq data (Fig 1C and Fig S5). Overall, scRNASeq data showed substantial bias toward immune populations (54% of cells) with underrepresentation of epithelial cells (2.3%), while snRNASeq had robust detection of epithelial cells (46%) and lesser detection of immune cells (15%). Comparison to a previous mouse 10x single cell dataset^8^ showed similar bias toward immune cells and reduced detection of mesenchymal cells (Fig 1D). Despite the reduced percentage of immune cells in snRNASeq data, clustering of immune cells in isolation resolved additional cell types, including classical and nonclassical monocytes, two populations of interstitial macrophages, NK cells and rare Il12b/Ccl22+ dendritic cells (Fig S2). Neutrophils and basophils were not detected in snRNASeq data. Finally, arterial, venous, capillary, and lymphatic endothelial cells were readily differentiated by snRNASeq (Fig S6).

### Merging of snRNASeq and scRNASeq data: co-clustering of mesenchymal cell types and differences in gene expression profiles

Because our scRNASeq and snRNASeq studies detected mesenchymal populations in similar proportion (Fig 1D), we selected these cells for further comparison. Count matrices from all snRNASeq and scRNASeq experiments were merged and re-clustered, with minimal batch effect after correction with the R package *Harmony* (Fig S3). Mesenchymal cells were then analyzed as a subset, with identification of six clusters, including three populations of matrix fibroblasts, myofibroblasts, pericytes, and smooth muscle cells (Fig 2A). All populations contained data from nuclei and cells (Fig 2B), further supporting the use of these cell types for gene expression comparisons between snRNASeq and scRNASeq. We next asked how gene detection differed between cells and nuclei. We found that a large majority of genes (96.5%) had less than 20% difference in expression between cells and nuclei. There were 329 genes (1.6%) that were detected in 25% more cells than nuclei, and 122 genes (0.6%) that were detected in 25% more nuclei than cells. Screening all detected genes (21,033) by log fold-change (>0.5, adjusted *P* value <0.05), revealed that only 3.0% (632) were enriched in cells, versus 1.4% (302) detected preferentially in nuclei (Fig 2D). We next performed gene ontology enrichment analysis on the cell vs. nucleus enriched genes. Unsurprisingly, genes associated with scRNASeq were related to ribosomal assembly and translation, as well as stress responses and apoptotic signaling, the latter potentially reflecting cell stress during dissociation (Fig 2E). snRNASeq-predominant genes were highly enriched for calcium transport and membrane depolarization, Slit/Robo signaling, and cAMP metabolism (Fig 2E).

**Figure 2.**
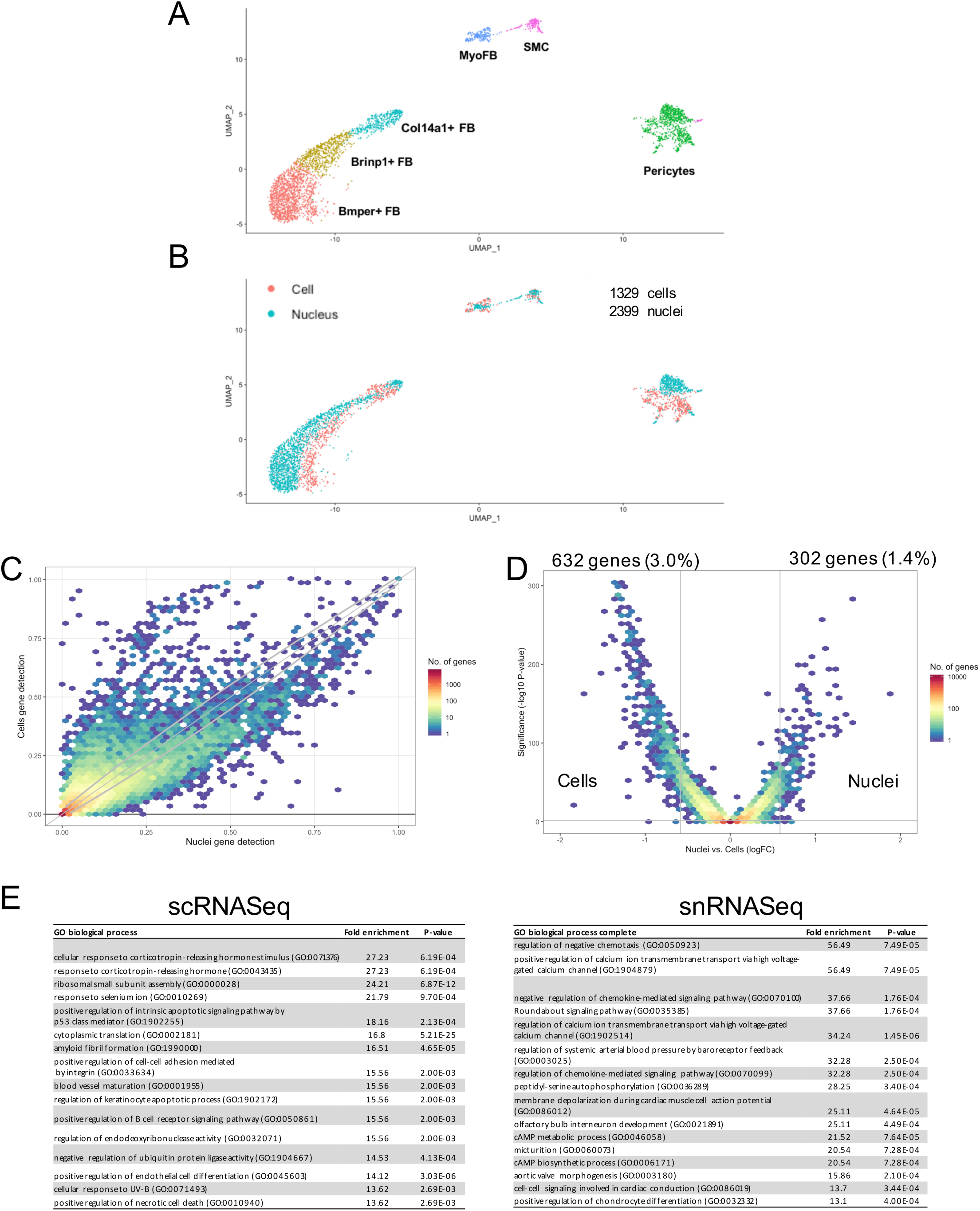
Merged single cell and single nucleus mesenchymal cells and differential expression between techniques. (A) uMAP plot of cell types after isolating mesenchymal subsets. (B) Overlap of transcriptomes from cells and nuclei after subclustering. (C) Binned scatterplot illustrating genes detected more reliably in cells versus nuclei, with grey lines marking the 95% confidence interval of variation detected by chance. (D) Volcano plot showing that 3.0% of genes are more highly expressed in cells (log fold change >0.5, adjusted *P* value<0.05), while 1.4% of genes are more highly expressed in nuclei. (E) Gene ontology analysis of enriched genes from scRNASeq and snRNASeq, arranged by log fold-change. FB, fibroblasts; MyoFB, myofibroblasts; SMC, smooth muscle cells; Peri, pericytes.

To further characterize gene detection differences between nuclei and cells, we examined expression of specific categories of genes. Among the genes predominantly sequenced in scRNASeq were those encoding mitochondrial, ribosomal, and heat shock proteins (Fig 3A). An advantage of snRNASeq over scRNASeq in other tissues has been reduced artifactual expression of stress-response genes, which are known to be induced during proteolysis at 37 C^14^. We could detect strong expression of the immediate early gene Fos in all cell types from the scRNA-seq dataset, but not from the snRNA-seq dataset (Fig. 3B). A panel of stress-induced genes including activator protein-1 (AP1) transcription factor component *Jun*, immediate early genes *Ier2* and *Ier3*, and stress sensor *Atf3* showed expression primarily in the single cell dataset (Fig 3C). In addition to background expression of stress-response genes, contamination from highly expressed genes in abundant cells (e.g. airway and alveolar epithelium) is a well-described phenomenon in single cell and single nucleus transcriptomics. While both mesenchymal populations from snRNASeq and scRNASeq showed contamination with epithelial genes, likely reflecting ambient mRNA, this was more significant in snRNASeq (Fig S3), contrary to previous results in kidney^7^. Genes more readily detected in nuclei included multiple transcription factors, as well as ion channels and signaling proteins (Fig 3D).

**Figure 3.**
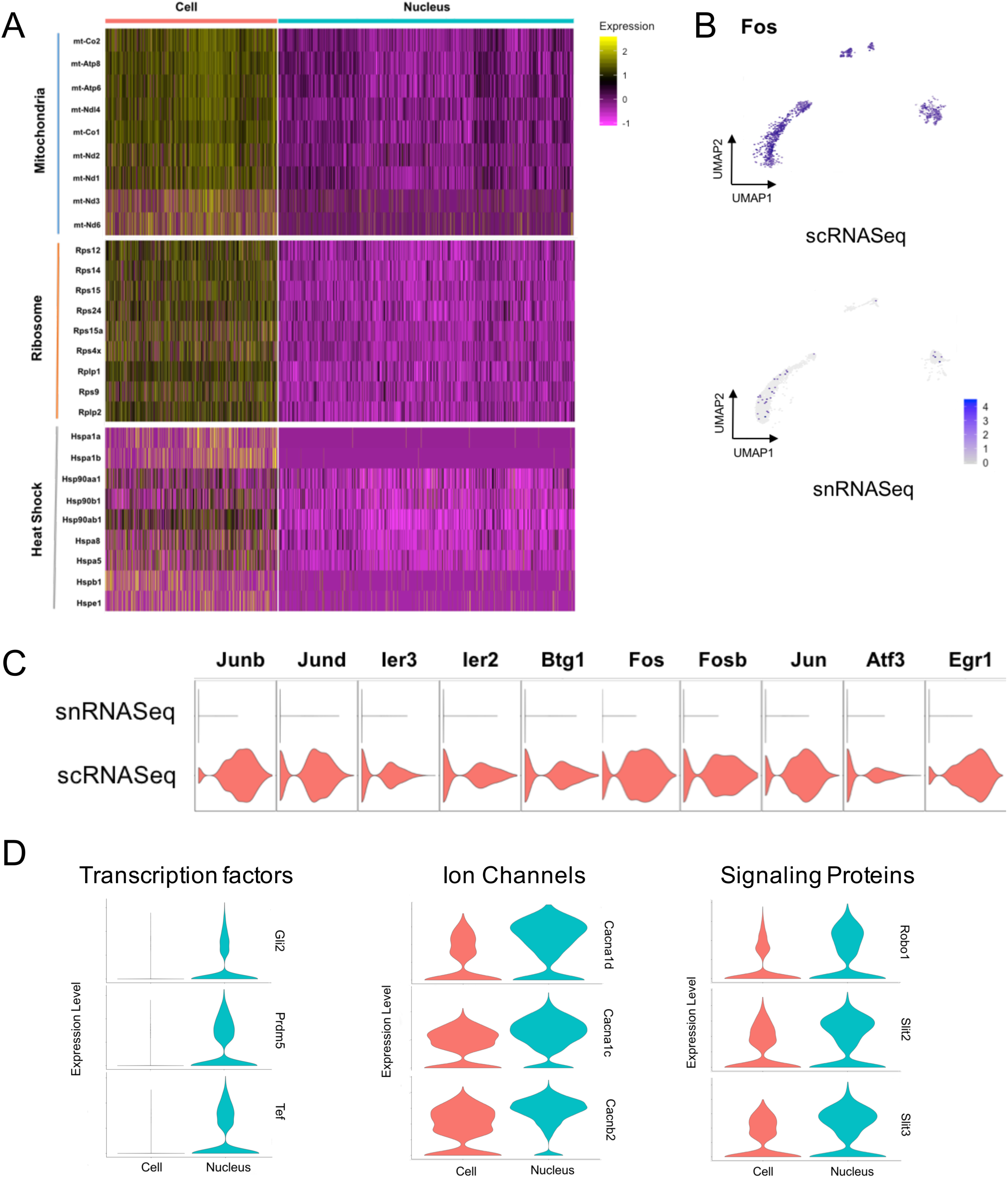
Differences in gene expression patterns between scRNASeq and snRNASeq. (A) Increased expression of mitochondrial, ribosomal, and heat shock response genes shown by heatmap. (B) Multiple stress response markers are enriched in single cell data, in a manner distributed equally among cell types (C). (D) Select genes with increased expression in snRNASeq data, including ion channels, signal transduction proteins, and transcription factors.

### Defining mesenchymal cell subpopulations

Previous lung single cell analyses in mice and humans have underrepresented fibroblasts and other mesenchymal populations^8,9^, which are embedded in matrix and less easily liberated during enzymatic tissue digestion. Given our robust recovery of mesenchymal cells in both scRNASeq and snRNASeq, we sought to further define the transcriptomes of these populations. Six populations were identified in our merged dataset (Fig 2A), including three populations of *Pdgfra*+ fibroblasts with distinct transcriptional profiles (Fig 4A and Fig 4B). While prior analyses have distinguished *Col14a1*- and *Col13a1*-expressing fibroblasts, the *Col13a1*+ population in our data contained two cell subtypes, the first characterized by expression of *Bmper, Fat3*, and *Fgfr4*, and the second by *Brinp1* and *Nalcn* (Fig 4B and Fig 4C). *Col14a1*+ fibroblasts additionally express *Dcn*, as well as *Lsamp* and transcription factor-encoding *Ebf2* (Fig 4C). Pericytes were defined by expression of the marker *Pdgfrb*, and expressed *Notch3, Pde5a*, and *Adcy8* (Fig 4C). Rarer populations of smooth muscle cells and myofibroblasts were also detected by snRNASeq and scRNASeq. Myofibroblasts were *Aspn, Grem2*, and *Hhip*+, and also expressed *Mapk4*, which encodes the atypical MAP kinase Erk4 and has not previously been described as a myofibroblast marker (Fig 4C). Smooth muscle cells were marked by *Acta2, Myh11*, and *Myocd* (Fig 4C).

**Figure 4.**
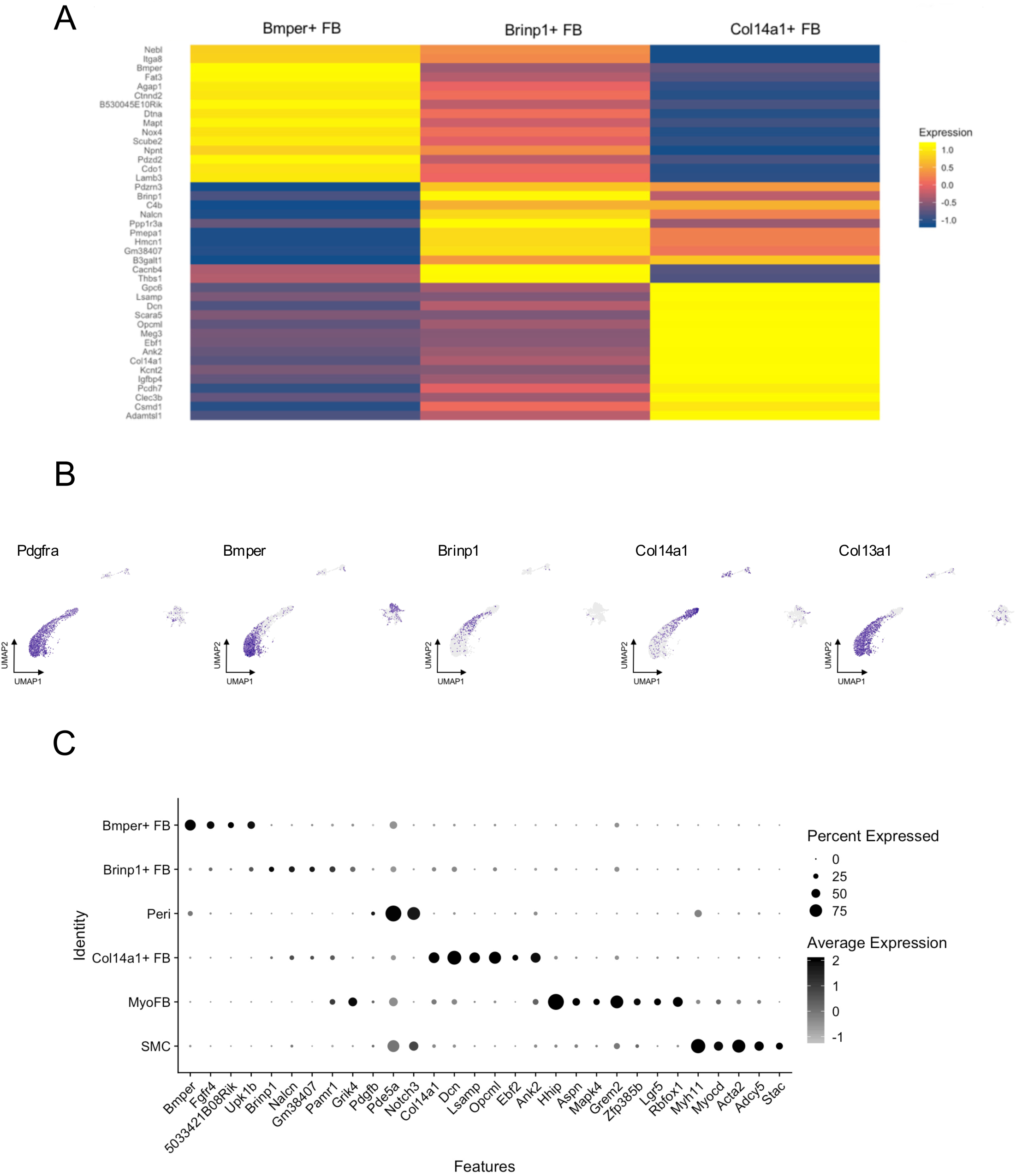
Characterization of mesenchymal cell types from combined scRNASeq and snRNASeq data. (A) Average gene expression heatmap showing distinct expression profiles of three fibroblast subtypes. (B) Of *Pdgfra+* fibroblasts, *Bmper*+ and *Brinp1*+ cells are *Col13a1*+ and do not express *Col14a1*. (C) Dot plot with additional marker genes for mesenchymal cell subtypes. FB, fibroblasts; MyoFB, myofibroblasts; SMC, smooth muscle cells; Peri, pericytes.

### Receptor-ligand interactome for the alveolar compartment based on snRNASeq data

Cell type-specific transcriptomic data in a tissue allows mapping of potential receptor-ligand interactions among cell types in close anatomic proximity. Because alveolar cell types were well represented in our snRNASeq data, we sought to define these signaling interactions in the alveolar compartment under control conditions by cross referencing our differentially expressed gene lists against an available receptor-ligand interaction database^15^. In the alveolus, AT1 cells encode a variety of signaling proteins, including *Vegfa, Bdnf, Wnt3a*, and *Pdgf*, whose corresponding receptors are found in endothelial cells, alveolar macrophages, and fibroblasts (Fig 5A, B), while additional *Pdgf* isoforms were produced by alveolar macrophages and capillary endothelial cells. Fibroblast interactions with epithelial cells include signaling through *Wnt5a* and *Igf1*.

**Figure 5.**
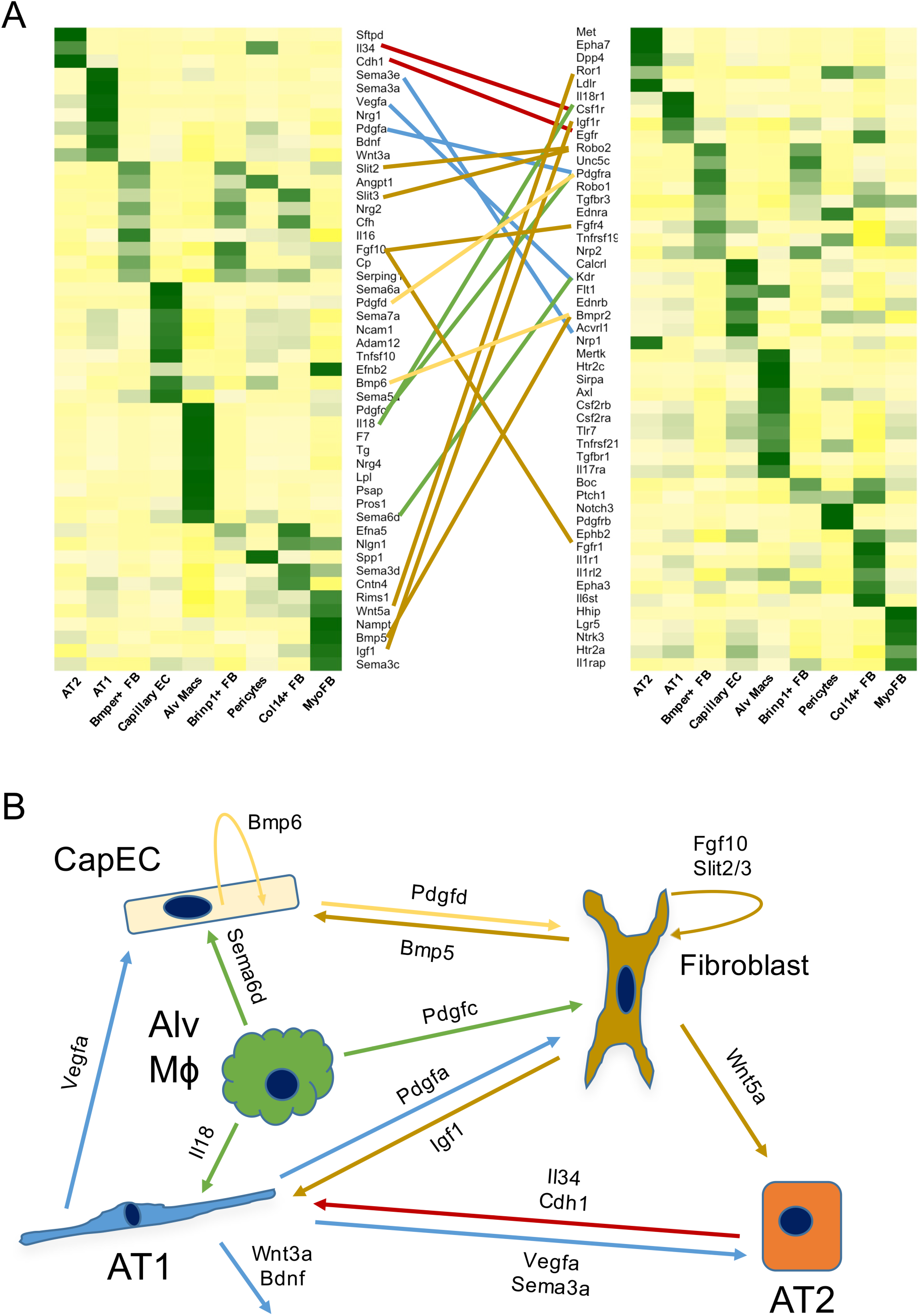
Ligand-receptor mapping in alveolar cell types. (A) Cell type-specific expression heatmaps of ligands and receptors illustrating potential networks of intercellular communication. (B) Alveolar signaling pathways suggested by the current snRNASeq data. AT1, alveolar type 1 cells; AT2, alveolar type 2 cells; FB, fibroblasts; MyoFB, myofibroblasts; EC, endothelial cells.

## Discussion

While scRNASeq has led to several important discoveries in lung, the application of this technology has practical limitations, perhaps the most important of which is the need for fresh tissue. The snRNASeq data in the present study was obtained from small pieces of snap frozen mouse lung, and could therefore be applied to existing banked samples. While we did not test this approach on tissue stored for months to years, studies evaluating RNA stability in cryogenically stored samples suggest that this may be a viable approach even after long-term storage^16^. We adapted our protocol from an existing method available for human lung nuclear isolation for snRNASeq^11^, eliminating the nuclear staining and flow sorting steps, which control for poor nuclear RNA quality. While nuclear RNA quality is difficult to assess, our experience in kidney suggests that generation of high quality snRNASeq data from cryopreserved samples is possible without flow sorting^17^ which simplifies the workflow substantially. Future efforts will focus on applying the current protocol to archived samples of normal and diseased human lung.

Inclusion of intronic and exonic reads when mapping snRNASeq data is known to improve gene detection, as confirmed in our data. Less expected was the effect of intron inclusion on scRNASeq data, which was still substantial though smaller in magnitude than for snRNASeq. Previous snRNASeq/scRNASeq comparisons have not observed this^6,7^. This phenomenon may relate to tissue- or cell type-specific differences in RNA processing, or to proprietary chemistry changes in newer 10x Genomics 3’v3 kits (e.g. harsher in-droplet lysis conditions leading to greater nuclear membrane disruption and release of nuclear mRNAs).

We observed a substantial bias toward detection of immune cell types in scRNASeq. The cell isolation protocol in the present study differs from previously published approaches for scRNASeq in its use of mechanical dissociation with GentleMacs and use of an Optiprep gradient to exclude RBCs and remove dead cells and degraded nucleotides (in lieu of RBC lysis and annexin V-based dead cell removal steps). While this may have contributed to the degree of observed dissociation bias, previous scRNASeq studies in mouse have shown similar propensities, underrepresenting alveolar and airway epithelial cells^8^. Notably, though mesenchymal cell detection has also been a weak point of prior scRNASeq studies, we observed similar numbers and subtypes of mesenchymal cells by snRNASeq and scRNASeq. Many epithelial populations including neuroendocrine cells and basal cells were seen only in snRNASeq. We did not detect ionocytes in our data, likely due to use of distal tissue samples (exclusion of trachea and main bronchi) and overall rarity.

A common issue encountered in single cell transcriptomics is amplification of contaminating background genes. In contrast to previous work, we found this problem to be more significant in single nucleus data, in which nearly all cells were found to express epithelial marker genes such as the club cell marker *Scgb1a1*. Further nuclear isolation protocol modifications (e.g. additional pellet washes) or computational approaches^18^ may help eliminate this contamination or limit its effect on downstream analysis. As previously reported, scRNASeq is also associated with “off-target” gene detection in the form of stress response/apoptotic genes and mitochondrial/ribosomal genes which may not be of interest in defining cell types or states^6,7^.

Previous studies have specifically explored the diversity of lung mesenchymal cells through lineage tracing and microarray analysis^19^, or more recently by scRNASeq^20^, generally selecting cells by flow cytometry prior to expression profiling. Through an unbiased whole-organ approach using snRNASeq we observed heterogeneity within *Pdgfra+*/*Col13a1*+ fibroblasts that had not been previously described. Where *Brinp1*+/*Col13a1*+ cells localize in the lung, and whether they represent a transition state between *Bmper*+ and *Col14a1*+ cells or a unique cell type remains to be defined. Additionally, resting myofibroblasts with relatively low *Acta2* and high *Hhip/Lgr5* expression were identified, similar to previous reports^20,21^. The presence of *Mapk4* in these cells is of note, given that its gene product Erk4 was recently implicated in cell proliferation and survival pathways through noncanonical mTOR signaling in cancer^22^, and that mTOR signaling has a well-described role in pulmonary fibrosis^23,24^. Defining the response of these cell types to injury and their degree of expansion in fibrosis will be a useful future application of snRNASeq.

Given the diversity of cell types and sequencing depth in snRNASeq, we interrogated the data for ligand-receptor interactions in the alveolar compartment. AT1 cells were identified as a robust source of signaling ligands in the alveolus at homeostasis. While best described during development, *Vegfa* signaling has also been associated with protection from acute lung injury^25^. Bdnf-Tkfb signaling has been implicated previously in epithelial-mesenchymal transition (EMT)^26^. While protein level data for cell type-specific expression are lacking, *Pdgfa, Vegfa, Wnt3a*, and *Bdnf* mRNA are all predominantly expressed in human AT1 cells in a large dataset including control, COPD, and IPF patients (IPFcellatlas.com)^3^. Of note, we also observed noncanonical Wnt pathway crosstalk between resting myofibroblasts and AT2 cells via *Wnt5a/Ror1*, and previous data suggest Wnt5a from adjacent fibroblasts maintains stem-ness in subsets of AT2 cells^27^. Fibroblasts were also a source of *Slit2*, whose loss of expression in fibrosis is reported to drive fibrocyte differentiation^28^. Similar analysis of snRNASeq data in disease conditions can provide insight into pathological intercellular crosstalk.

In conclusion, snRNA-seq is feasible from cryopreserved lung, and our simplified protocol eliminates the need for FACS purification. In comparison to scRNA-seq, our protocol offers equivalent gene detection, eliminates artifactual transcriptional stress responses and delivers a much higher proportion of epithelial cells.

## Acknowledgements

Work in the Humphreys Lab is supported by NIH/NIDDK grants DK103740 and DK107374 (all to B.D.H).

