## Supplemental Figures 1-6 for "Single nucleus RNASeq profiling of mouse lung: reduced dissociation bias and improved detection of rare cell types compared with single cell RNASeq"

This PDF file includes: Figures S1-S6

### Figure S1

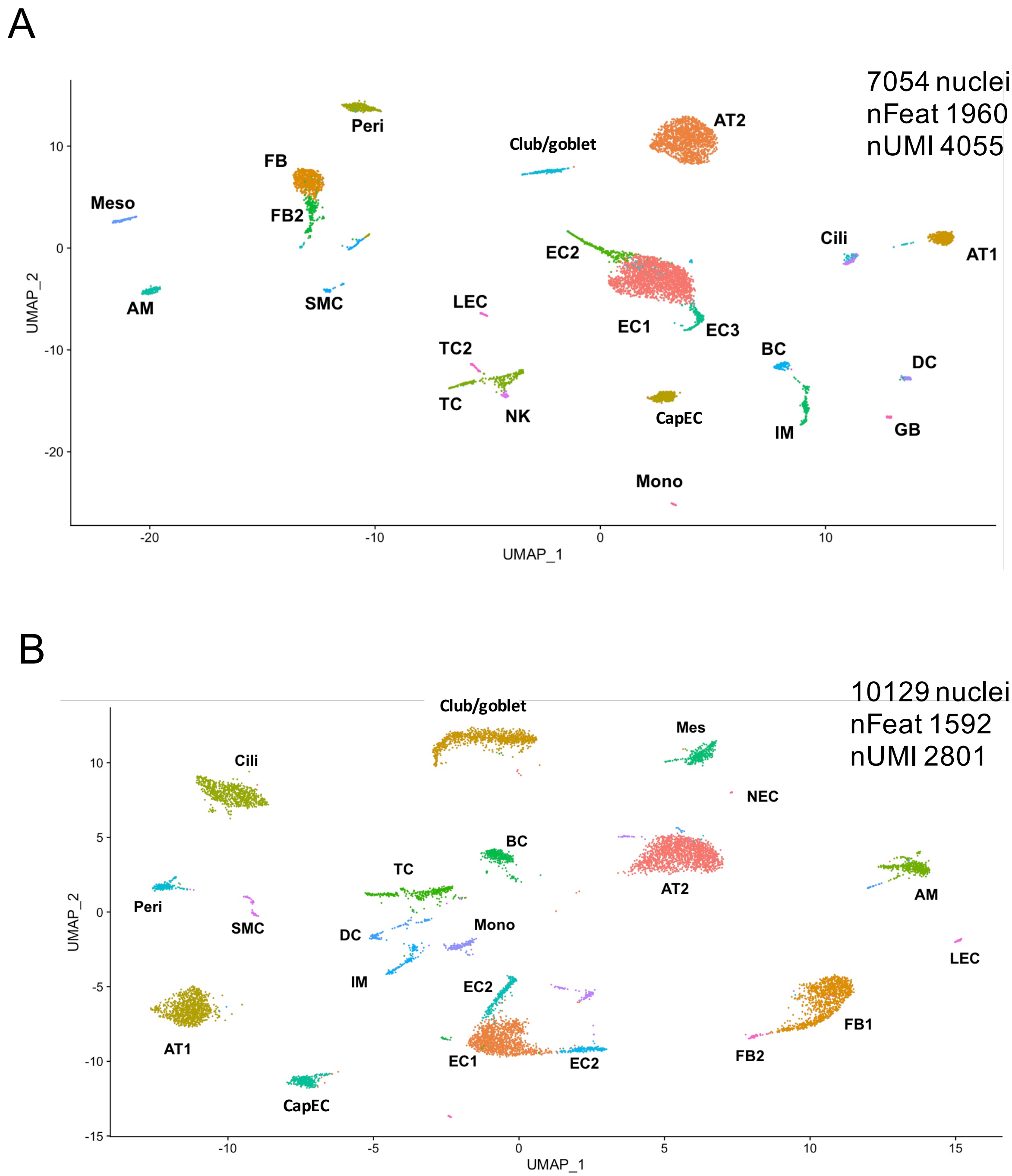


**Figure S1. uMAPs for individual snRNASeq studies on control mouse lung.** Overall similar gene and cell type detection between experiments, with the first (A) improved in terms of genes and UMI detected per nucleus. Neuroendocrine cells were identified in one of two runs (B). EC, endothelial cells; LEC, lymphatic endothelial cells; AT1, alveolar type 1 epithelial cells; AT2, alveolar type 2 epithelial cells; BC, B cells; GB, germinal B cells; TC, T cells; DC, dendritic cells; FB, fibroblasts; AM, alveolar macrophages; IM, interstitial macrophages; Peri, pericytes; Mono, monocytes; NK, natural killer cells; Mes, mesothelial cells; SMC, smooth muscle cells; NEC, neuroendocrine cells; nFeat, number of features per nucleus; nUMI, number of UMI per nucleus.

### Figure S2

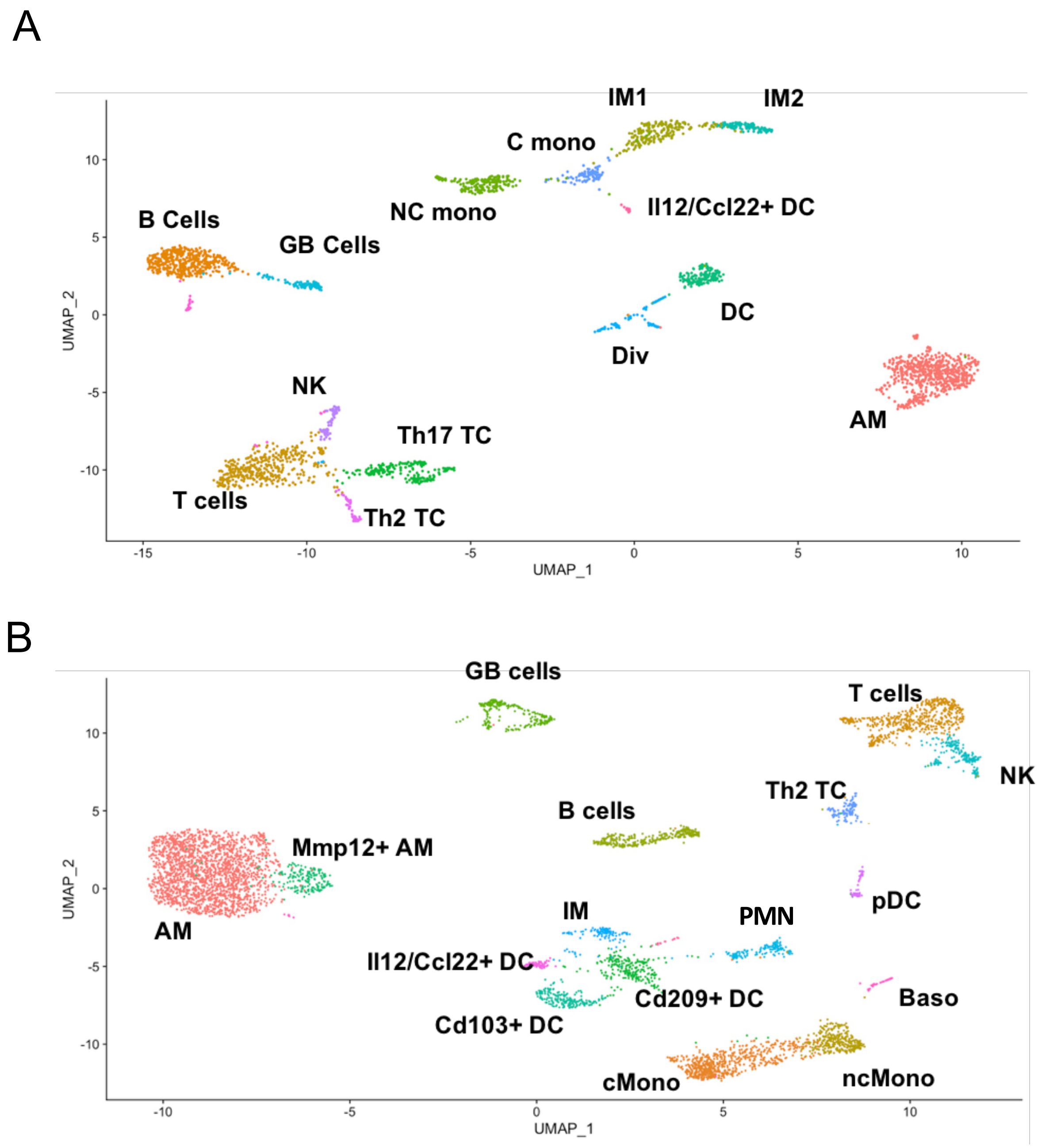


**Figure S2. Immune cell sub-clustering in snRNASeq and scRNASeq.** (A) uMAP of immune populations from combined single nucleus data, with 14 cell types including subpopulations of interstitial macrophages and Th17 T cells. (B) Immune populations from scRNASeq, with improved resolution of dendritic cell subtypes, neutrophils, and basophils. AM, alveolar macrophages; IM, interstitial macrophages; DC, dendritic cells; cMono, classical monocytes; ncMono, nonclassical monocytes; Baso, basophils; PMN, neutrophils; pDC, plasmacytoid dendritic cells; NK, NK cells; GB, germinal B cells; TC, T cells; Th2, T helper 2; Th17, T helper 17.

### Figure S3

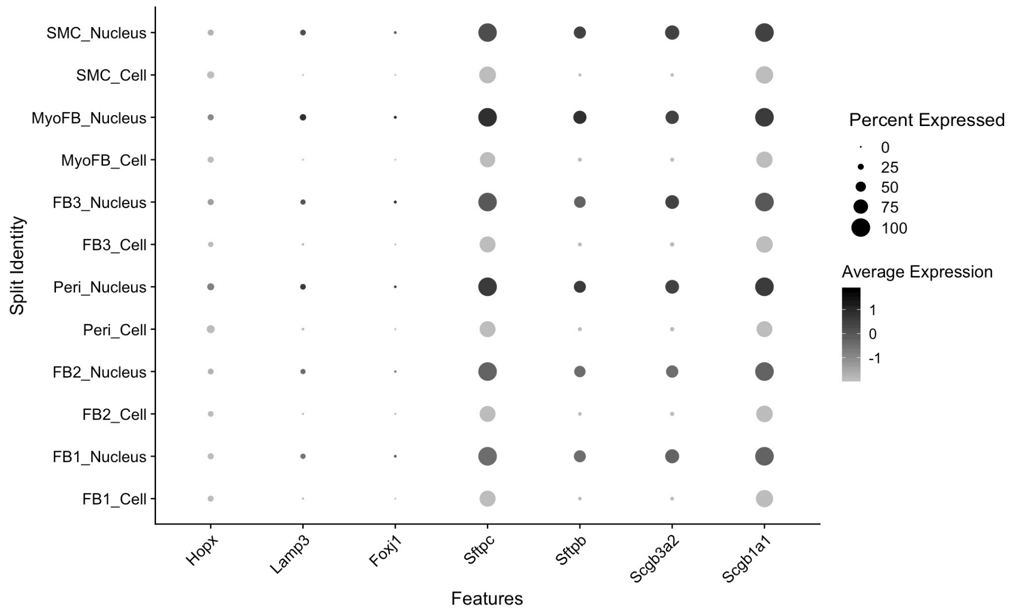


**Figure S3. Contamination of mesenchymal cell transcriptomes by epithelial genes in snRNASeq versus scRNASeq.** Dot plot showing diffuse detection of alveolar and airway epithelial genes, particularly club cell and alveolar type 2 cell markers, in single nucleus data more than single cell data. SMC, smooth muscle cells; MyoFB, myofibroblasts; FB, fibroblasts; Peri, pericytes.

### Figure S4

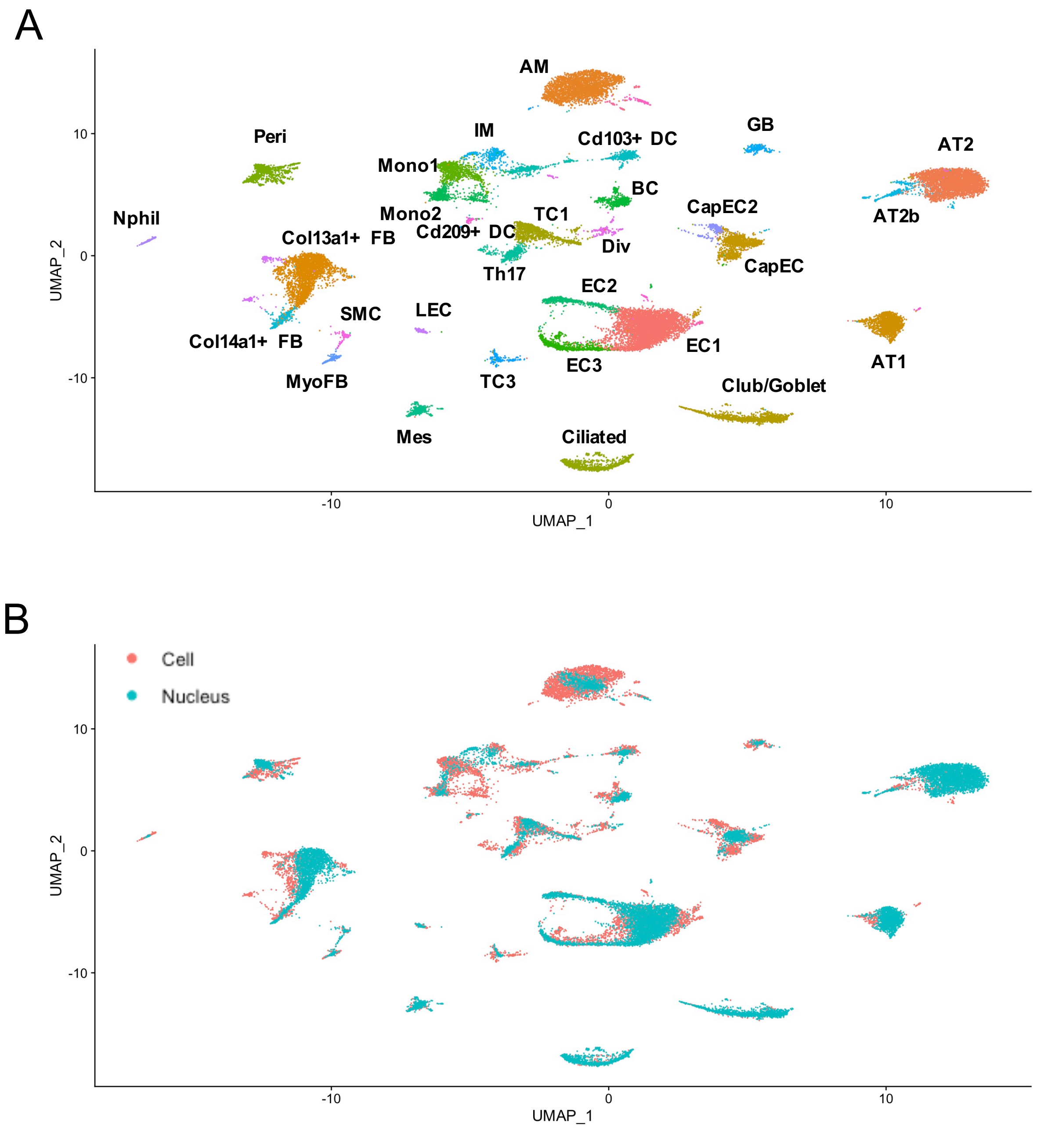


**Figure S4. Merged single cell and single nucleus datasets.** (A) uMAP of scRNASeq and snRNASeq data after merger and batch correction, showing 30 distinct cell types. (B) Separately labeled single cell and nucleus transcriptomes shows minimal batch effect after correction with *Harmony*. EC, endothelial cells; LEC, lymphatic endothelial cells; AT1, alveolar type 1 epithelial cells; AT2, alveolar type 2 epithelial cells; BC, B cells; GB, germinal B cells; TC, T cells; DC, dendritic cells; FB, fibroblasts; MyoFB, myofibroblasts; AM, alveolar macrophages; IM, interstitial macrophages; Peri, pericytes; Mono, monocytes; NK, natural killer cells; Mes, mesothelial cells; SMC, smooth muscle cells.

### Figure S5

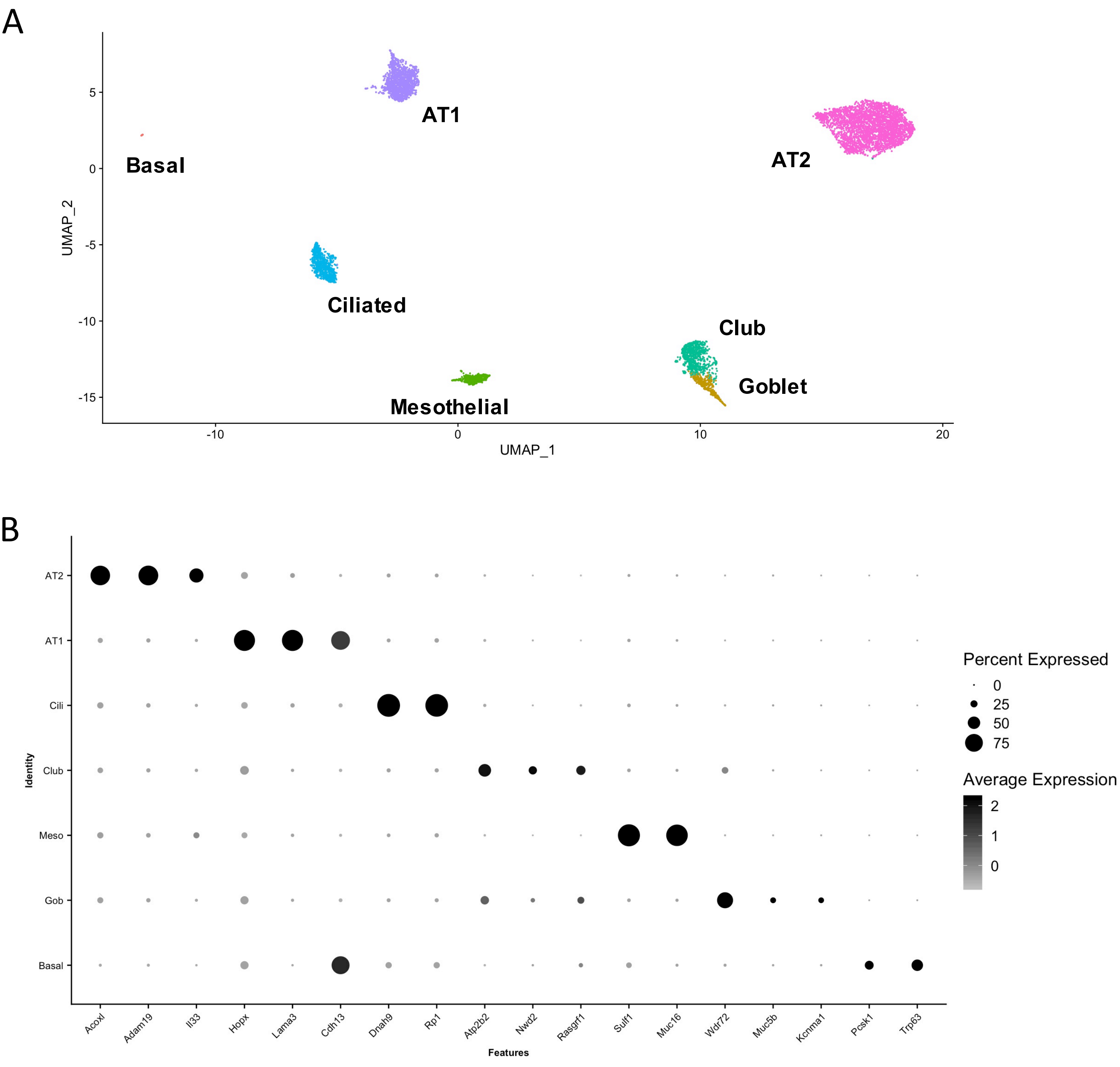


**Figure S5. Epithelial subpopulations in snRNASeq data.** (A) uMAP of epithelial cell types from combined snRNASeq data including basal and goblet cells. (B) Marker genes for each of the epithelial clusters. AT1, alveolar type 1; AT2, alveolar type 2; Cili, ciliated; Meso, mesothelial; Gob, goblet.

### Figure S6

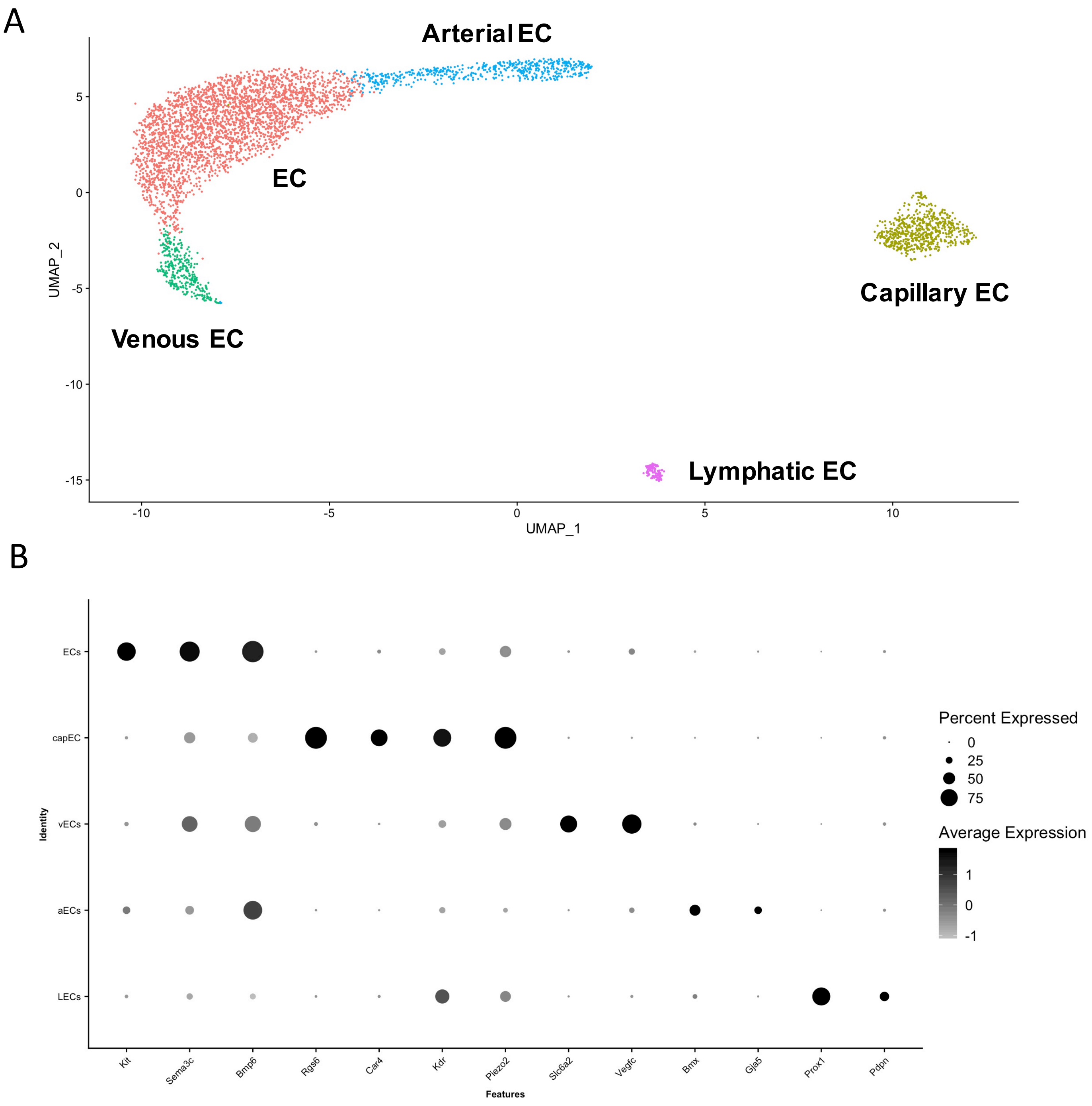


**Figure S6. Endothelial subpopulations in snRNASeq data.** (A) uMAP of endothelial cell types from combined snRNASeq data including distinct arterial, venous, lymphatic, and capillary endothelial cells (EC). (B) Dot plot of marker genes for subtypes of endothelial cells. capEC, capillary endothelial cells; vECs, venous endothelial cells; aECs, arterial endothelial cells; LECs, lymphatic endothelial cells.
